# Epigenome-wide association analysis of daytime sleepiness in the Multi-Ethnic Study of Atherosclerosis reveals African-American specific associations

**DOI:** 10.1101/447474

**Authors:** Richard Barfield, Heming Wang, Yongmei Liu, Jennifer A Brody, Brenton Swenson, Ruitong Li, Traci M. Bartz, Nona Sotoodehnia, Yii-der I. Chen, Brian E. Cade, Han Chen, Sanjay R. Patel, Xiaofeng Zhu, Sina A. Gharib, W. Craig Johnson, Jerome I. Rotter, Richa Saxena, Shaun Purcell, Xihong Lin, Susan Redline, Tamar Sofer

## Abstract

**Study Objectives:** Excessive daytime sleepiness (EDS) is a consequence of inadequate sleep, or of a primary disorder of sleep-wake control. Population variability in prevalence of EDS and susceptibility to EDS are likely due to genetic and biological factors as well as social and environmental influences. Epigenetic modifications (such as DNA methylation-DNAm) are potential influences on a range of health outcomes. Here, we explored the association between DNAm and daytime sleepiness quantified by the Epworth Sleepiness Scale (ESS).

**Methods:** We performed multi-ethnic and ethnic-specific epigenome-wide association studies for DNAm and ESS in 619 individuals from the Multi-Ethnic Study of Atherosclerosis. Replication was assessed in the Cardiovascular Health Study (CHS). Genetic variants in genes proximal to ESS-associated DNAm were analyzed to identify methylation quantitative trait loci and followed with replication of genotype-sleepiness associations in the UK Biobank.

**Results:** 61 methylation sites were associated with ESS (FDR ≤ 0.1) in African Americans only, including an association in *KCTD5*, a gene strongly implicated in sleep. One association (cg26130090) replicated in CHS African Americans (p-value 0.0004). We identified a sleepiness-associated methylation site in the gene *RAI1*, a gene associated with sleep and circadian phenotypes. In a follow-up analysis, a genetic variant within *RAI1* associated with both DNAm and sleepiness score. The variant’s association with sleepiness was replicated in the UK Biobank.

**Conclusions:** Our analysis identified methylation sites in multiple genes that may be implicated in EDS. These sleepiness-methylation associations were specific to African Americans. Future work is needed to identify mechanisms driving ancestry-specific methylation effects.

**Statement of Significance:** Excessive daytime sleepiness is associated with negative health outcomes such as reduction in quality of life, increased workplace accidents, and cardiovascular mortality. There are race/ethnic disparities in excessive daytime sleepiness, however, the environmental and biological mechanisms for these differences are not yet understood. We performed an association analysis of DNA methylation, measured in monocytes, and daytime sleepiness within a racially diverse study population. We detected numerous DNA methylation markers associated with daytime sleepiness in African Americans, but not in European and Hispanic Americans. Future work is required to elucidate the pathways between DNA methylation, sleepiness, and related behavioral/environmental exposures.

## Introduction

Excessive daytime sleepiness (EDS), is estimated to affect between 10 to 20% of the population^1,2^. EDS is associated with numerous adverse clinical, behavioral and public health outcomes, including work and vehicular accidents^3-10^; reduced health-related quality of life^11-13^; cognitive and performance deficits^14,15^; and increased rates of stroke and total and cardiovascular mortality^16,17^. There are multiple mechanisms for EDS, including insufficient sleep duration due to behavioral, social or work-related factors; sleep disruption due to a sleep disorder (sleep apnea, periodic limb movement disorder); circadian misalignment; or the presence of a primary disorder of hypersomnia that affects the central sleep-wake control processes (e.g., narcolepsy). The prevalence of EDS has increased over the past decades, likely due to an increase in working hours and availability of electronic devices^3^. A rise in EDS prevalence may also reflect the increased prevalence of obesity, which is associated with increased sleepiness^18,19^. This association is hypothesized to be due to obesity-related co-morbidities that reduce sleep quality, including obstructive sleep apnea, as well as to metabolic and neuroendocrine effects of adipokines on wake promoting neurons^20^.

Although EDS is prevalent in the population, there appears to be large inter-individual differences in propensity for sleepiness following sleep deprivation, sleep fragmentation, or sleep apnea^21,22^. Similarly, there is significant population variability in prevalence of EDS^23^. Differences in sleepiness, including vulnerability or resilience to sleep disrupting influences, have been suggested to be due to genetic and other biological differences, although social and environmental influences also likely play a role^24,25^. Generally, the bases for differences in sleepiness between populations are not well understood.

Epigenetic modifications are increasingly recognized to mediate the impact of environmental influences on gene expression and on prevalence and severity of a wide range of health outcomes, including neuropsychiatric, metabolic and cardiovascular diseases. The most studied epigenetic marker, DNA methylation (DNAm), occurs when a methyl group is added to a cytosine that is followed by a guanine on the genome (a “CpG” site). Changes in DNA methylation occur in response to a wide range of exposures, many of which are associated with sleep and sleepiness, such as obesity, diet, and stress^26-32^. Therefore, we postulated that changes in DNAm would associate with variation in sleepiness and these changes may be population-specific, providing insights into underlying susceptibility to sleepiness that may be partially attributed to environmental factors. To the best of our knowledge, there has been no reported study on DNA methylation and sleepiness.

In this paper, we performed an Epigenome Wide Association Study (EWAS) of EDS quantified by the Epworth Sleepiness Scale (ESS) in the Multi Ethnic Study of Atherosclerosis. We leveraged the racial/ethnic diversity of the sample to explore potential differences in EWAS associations by background groups. We performed replication analysis of methylation sites-ESS associations in the Cardiovascular Health Study (CHS) and identified cis-meQTLs in genes harboring associated methylation sites for replication analysis in the UK Biobank.

## Methods

### Study Sample

The study population consisted of participants from the Multi-Ethnic Study of Atherosclerosis (MESA), a prospective, longitudinal cohort study established to study factors associated with the development of cardiovascular disease. MESA clinic visits were first performed between 2000 and 2002 in six field centers across the United States when participants were free of known cardiovascular disease^33^. The subset of MESA individuals in this study is composed of those who participated in a sleep examination conducted in conjunction with MESA Exam 5 (described in detail previously^23,33^), and who also participated in the MESA DNAm study^34^. The blood draws for the methylation study were obtained during MESA Exam 5 (2010-2012)^35^ on a random subset of MESA participants at four of the six field centers: John Hopkins University, University of Minnesota, Columbia University, and Wake Forest University. A total of 623 individuals were available with both sleep data and DNAm data with four removed due to having missing ESS. The final study sample included 619 individuals: 132 African Americans (AAs), 202 Hispanic Americans (HAs), and 285 European Americans (EAs). The study was approved by the Institutional Review Boards of each participating site, and participants provided written informed consent, including use of genetic data.

### DNAm Collection and Processing

Methods for collection and assays for DNAm have been described previously^35^. In brief, peripheral blood was separated into mononuclear cells (CD14+) within two hours of collection. DNAm in monocytes was measured using the Illumina HumanMethylation450 BeadChip. Residual cell contamination in the monocyte data was assessed using Gene Set Enrichment Analysis, providing enrichment scores for neutrophils, B cells, T cells, and Natural Killer cells from Gene Expression data collected using the Illumina HumanHT-12 v4 Expression BeadChip and Illumina Bead Array Reader^35,36^. The DNAm data underwent quality control tests prior to analysis using the lumi Bioconductor package^37^. These included color bias adjustments via smooth quantile normalization, median background adjustment, standard quantile adjustment, checks for potential sex or race mismatches, and outlier detection via multidimensional plots. Additional details on pre-processing of the DNAm data can be found in Liu et al^34^. Following DNAm preprocessing, each of 484,882 methylation probes for each person has a Beta-value, representing the proportion of methylated monocytes at that site and person.

We further prepared the methylation data for analysis based on meta information and sample-specific characteristics, as follows. First, we excluded 61,219 probes that were within 10bp of a SNP with a minor allele frequency greater than 1% in European, African, or American populations from the 1000 genomes reference data^38^. This was done to exclude loci where variation is solely due to genetic polymorphisms. We also removed any non-cg methylation probes and probes on sex chromosomes. Next, we focused on 73,082 CpG sites that had an interdecile range greater than 0.1. Methylation values in these sites are unlikely driven by technical variability and focusing on them will reduce the multiple testing burden^39^. Then, we used the software program ComBat^40^ on these 73,082 CpG sites to remove any signal due to technical artifacts of chip and position on chip effects, while maintaining correlation with our primary set of covariates (self-reported ancestry, recruitment site, sex, age, and residual cell type enrichment). ComBat was run on the M-values (logit transformed Beta-values)^41^. We then removed any cross-reactive probes as defined by Chen et al^42^, leaving us with 66,028 DNAm probes to analyze.

### Single Nucleotide Polymorphism (SNP) Data

For consenting individuals, DNA was extracted from whole blood and genotyped on Affymetrix 6.0 GWAS array. Standard quality control methods for SNP- and sample-level quality were applied, including the exclusion of participants and SNPs with over 5% missing call rates. This resulted in 895,289 genotyped variants in 615 individuals with DNAm data. For downstream analysis, we excluded SNPs with minor allele frequency (MAF) less than 1%. Further details on the genotype and quality control can be found in Vargas et al^43^. We calculated the top 5 principal components (PCs) from the Linkage-Disequilibrium pruned set SNPs with MAF≥5%. PCs were calculated in the combined sample and also in each ethnic-specific group. There were four AA individuals that had missing SNP information; therefore, the PCs were imputed with the mean value of the respective PC in the AA subset that had SNP information.

### Sleep Assessments

As part of the MESA Sleep Examination (2010-2013), participants completed standardized questionnaires and underwent a single night in-home polysomnography (Compumedics Somte Systems, Abbotsville, Australia, AU0) and 7 day wrist actigraphy (Philips-Respironics Spectrum, Murrysville, PA), as described before^23^. The primary sleep measure was daytime sleepiness as quantified by the Epworth Sleepiness Scale (ESS), an 8-item validated instrument that asks the individual to assess likelihood of dozing off in a variety of daily activities using a 4 point (0-3) scale^44^. ESS scores vary from 0 to 24, with higher scores denoting more sleepiness^44^. ESS was assessed within one year of blood draw for DNA methylation. We identified a set of additional sleep phenotypes/exposures that might be associated with both DNAm and ESS, and thus are potential mediators or confounders of any DNAm and ESS association. These measures were: insomnia (report of doctor diagnosed insomnia or the Women’s Health Initiative (WHI) Insomnia Rating Scale^45^, a validated scale varying from 0 to 20); Apnea-Hypopnea Index [AHI] (assessed by polysomnography as the sum of all apneas and hypopneas associated with 3% or more oxygen desaturation divided by total sleep time); overnight hypoxemia (derived by polysomnography as the percentage of sleep time the participant spent with oxyhemoglobin saturation less than 90% [Per90]); and sleep duration (actigraphy derived average sleep duration over the 7 monitoring nights as described before^23^).

### Covariates

Age, sex and race/ethnicity were self-reported. Other behavioral, socioeconomic and lifestyle exposures that may confound sleepiness-methylation associations were also assessed. Alcohol use was based on self-reported (yes/no); smoking status was classified by ever/former/never); depressive symptoms were based on responses to the Center for Epidemiological Studies (CES) depression scale^46^; having less than a college education; mother having less than a college education, the latter two variables representing socio-economic status. Body mass index (BMI) was calculated from weight and height measured at MESA Exam 5. Dietary variables were derived from a food frequency questionnaire: long chain score (sum of Omega fatty acids mg/d), total fats, carbohydrates, and saturated fatty acids (rams) and Total Alternative Healthy Eating Index - 2010 (Total AHEI-2010)^47^.

### Statistical Analysis

Our complete analysis approach is depicted in Figure 1. We first describe our discovery analyses, followed by sensitivity, gene expression, local methylation expressive quantitative trait loci (cis-meQTL, and replication analyses.

**Figure 1:**
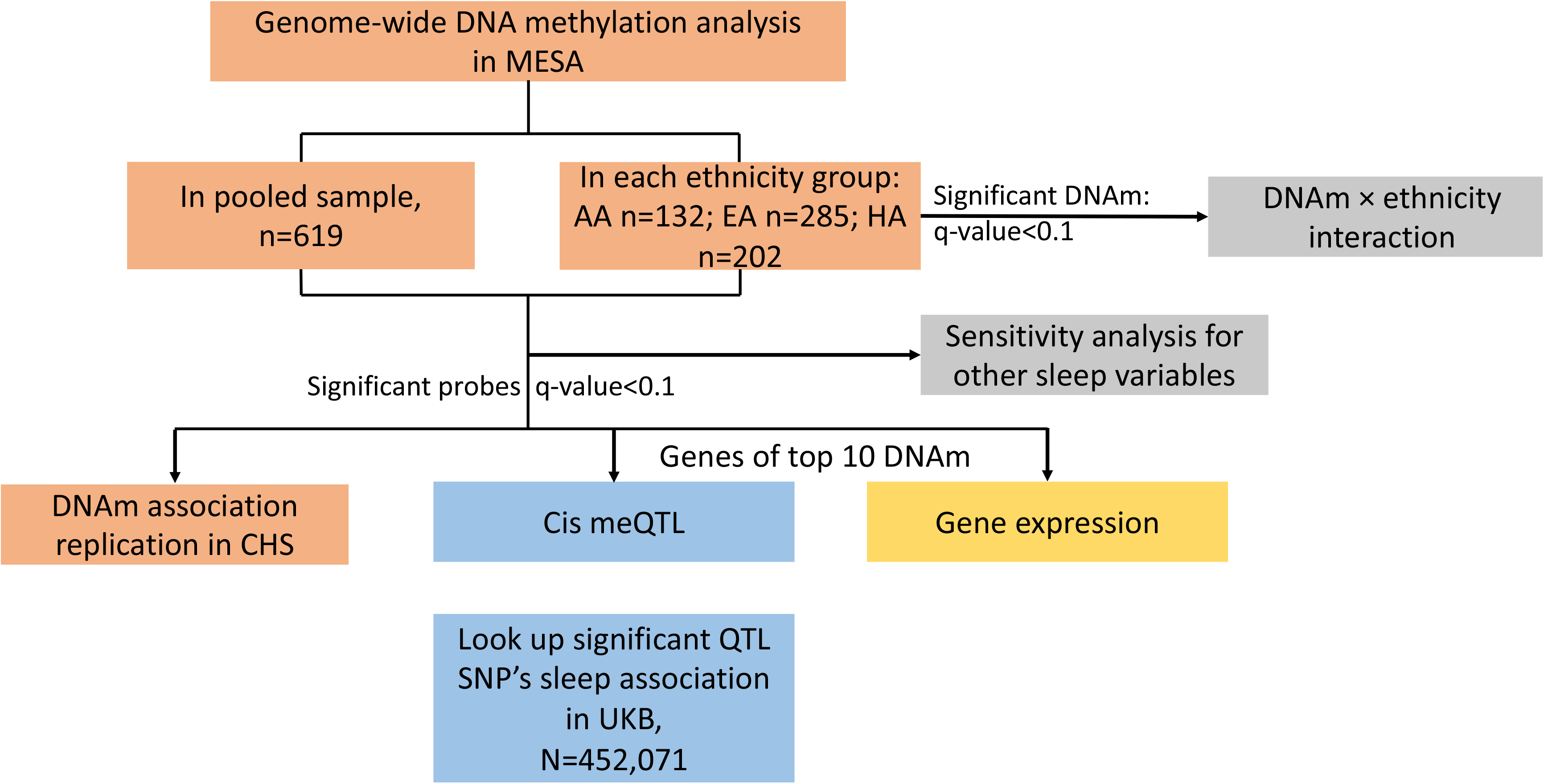
Analysis workflow for assessing associations between DNAm and ESS.

### Discovery Analysis for Methylation-ESS Association

The primary analysis outcome was ESS score, square-root transformed to achieve approximate normality. It was analyzed in a linear regression on DNAm M-values generated following application of ComBat (as described earlier). The primary analyses were adjusted for sex, age, residual cell type enrichment, recruitment site, and the top 5 genetic PCs. We first analyzed the transformed ESS on the combined multi-ethnic sample with an additional covariate for ethnicity (EA, AA, or HA) and PCs calculated in the overall group. We then performed race/ethnic group-specific analyses adjusting for the top five PCs calculated exclusively for each respective race/ethnicity group. To account for multiple testing, we calculated false discovery rate (FDR) controlling q-values in each of the analyses, i.e. the multi-ethnic and each ethnic-group specific analyses^48^. An association was deemed as significant if the q-value was less than 0.1^48^, thus controlling the FDR at the 10% level in each of the analyses. We evaluated heterogeneity in any race/ethnic specific results by fitting a linear model using the overall sample with self-reported race/ethnicity interaction terms with DNAm and testing these interaction terms using an F-test via analysis of variance (ANOVA).

### Sensitivity Analysis for methylation-ESS associations

We next identified behavioral, socioeconomic and lifestyle exposures, including variation in sleep and sleep disturbances, that may influence the sleepiness-methylation associations and further adjusted for these factors in sensitivity analyses conducted in significant probe-trait primary associations. Covariates considered for the sensitivity analysis were: BMI, sleep duration (actigraphy-based), insomnia, AHI, nocturnal hypoxemia [Per90], alcohol use, smoking status, CES depression scale, having less than a college education, mother having less than a college education, and dietary variables, as defined earlier in the manuscript. We also examined moderate sleep apnea defined as an AHI greater than 15, less than five hours of sleep, and overnight hypoxemia (dichotomized as Per90 greater than 5%). We only assessed variables that had a p-value less than 0.1 for their association with ESS after adjusting for age and sex. These were included together as covariates in the regression model for CpG sites with a q-value less than 0.1. We handled missingness of continuous variables (up to five missing values per variable) by imputing the missing value to the mean of the variable calculated in the rest of the study participants of the same self-reported race, and removed individuals with missing categorical values. Missing information by variable is provided in Supplementary Table 1. Finally, we evaluated whether an observed change in effect size estimate between a methylation site and ESS after including each or sets of covariates was higher than by chance via permutation. We permuted the additional exposure/sensitivity variables 10,000 times and calculated the proportion of permutations in which the observed change in DNAm-ESS effect sizes was higher compared to the DNAm-ESS effect size in the un-permuted sensitivity variables.

### DNAm-ESS replication analysis in the Cardiovascular Health Study (CHS)

The CHS is a population-based cohort study of risk factors for coronary heart disease and stroke in adults ≥65 years conducted across four field centers^49^. The original predominantly European ancestry cohort of 5,201 persons was recruited in 1989-1990 from random samples of the Medicare eligibility lists; subsequently, an additional predominantly African-American cohort of 687 persons was enrolled for a total sample of 5,888. CHS was approved by institutional review committees at each field center and individuals in the present analysis had available DNA and gave appropriate informed consent for use of genetic information. DNA methylation was measured on 336 European ancestry and 329 African-American ancestry participants at study year 5. The samples were randomly selected among participants without presence of coronary heart disease, congestive heart failure, peripheral vascular disease, valvular heart disease, stroke or transient ischemic attack at study baseline or lack of available DNA at study year 5. Sleepiness was assessed at year 6 using the ESS.

Methylation measurements were performed at the Institute for Translational Genomics and Population Sciences at the Harbor-UCLA Medical Center--Los Angeles Biomedical Research Institute using the Infinium HumanMethylation450 BeadChip (Illumina Inc, San Diego, CA). Quality control was performed in in the minfi R package^50-52^ (version 1.12.0, http://www.bioconductor.org/packages/release/bioc/html/minfi.html). Samples with low median intensities of below 10.5 (log2) across the methylated and unmethylated channels, samples with a proportion of probes falling detection of greater than 0.5%, samples with QC probes falling greater than 3 standard deviation from the mean, sex-check mismatches, failed concordance with prior genotyping or > 0.5% of probes with a detection p-value > 0.01 were removed. In total, 11 samples were removed for sample QC resulting in a sample of 323 European-ancestry and 326 African-American samples. Methylation values were normalized using the SWAN quantile normalization method^52^. Since white blood cell proportions were not directly measured in CHS they were estimated from the methylation data using the Houseman method^53^.

We performed replication look-up of the significant (q-value<0.1) associations in CHS AA and EA. We adjusted for age, sex, recruitment site, current smoking status, white blood cell counts, and top five genetic PCs. For this, we examined results in both individual methylation sites, and used a binomial test^54^ for the overall agreement in the direction of associations between discovery and replication studies. A methylation site was considered to be statistically associated with ESS if its one-sided p-values (with direction of association guided by the direction of DNAm-sleepiness association in MESA discovery) was less than 0.05 divided by number of FDR significant sites in MESA, a Bonferroni threshold controlling the Family-Wise Error Rate of replication^55^. We analyzed both CHS AA and EA samples separately.

### Testing for cis-meQTLs in MESA

We hypothesized that a combination of DNA methylation and genetic association plays a role in sleepiness. To study this hypothesis, we attempted to identify variants that are associated with both methylation and sleepiness. From a causal pathway perspective, this analysis considers the setting in which a genetic variant predicts both DNAm and sleepiness, where the association with sleepiness is either direct (pleiotropic) or mediated through DNAm. An example of the latter mechanism, a genetic variant could affect the methylation status of a CpG site by disrupting binding sites for regulatory proteins, which in turn influences sleepiness via a molecular mechanism not directly related to the product of the gene of interest. Due to our limited sample size, we implemented a screening process on genetic variants in the CpG region, followed by a replication analysis of the SNP-sleepiness association in an independent cohort (UK Biobank).

First, we mapped the top ten methylation sites to their respective genes to assess if genetic variants in that gene were associated with both ESS and DNAm. Specifically, for each gene that a top probe mapped to, we first performed association analysis between genotyped SNPs within 100kb of the gene and the associated DNAm site adjusting for age, sex, top five PCs, study site, and residual cell type. This identified local (cis) meQTLs (screening step 1). SNPs with associations significant at P<0.005 (accounting for ten comparisons) were next assessed for their association with sleepiness, with and without additional adjustment for the DNAm (screening step 2). SNPs with P<0.05 in screening step 2 were carried forward to independent replication analysis (see below) of the SNP-sleepiness association in the UK Biobank.

### Replication analysis of cis-meQTL-sleepiness association in the UK Biobank

We further looked up the SNP-sleepiness associations using individuals of European ancestries from a GWAS of sleepiness in UK Biobank. The UK Biobank is a large prospective study for a wide range of genetics and health outcomes in over 500,000 participants aged 40-69 years recruited from 2006 to 2010 in the United Kingdom^56^. Genotyping and imputation data are described in Bycroft et al. (20 1 7)^57^. Sleepiness was assessed by self-reported responses to the question “How likely are you to dose off or fall asleep during the daytime when you don’t mean to? (e.g. when working, reading or driving)” with the options of “Never/rarely”, “sometimes”“often”, “all of the time”, “do not know”, and “prefer not to answer.” Association analysis was performed in 452,071 individuals adjusted for age, sex, genotyping array, 10 PCs and genetic relatedness matrix.

### Gene expression analysis

For genes corresponding to the top 10 DNAm sites, we used gene expression data in MESA to study the evidence for association between DNAm and gene expression (of the same gene), and gene expression association with ESS. Gene expression profiling and processing are described in detail in Liu at al^34^. In brief, gene expression was measured using Illumina HumanHT-12 v4 Expression BeadChip array in mononuclear cells, on the same participants with DNA methylation measures. We performed (1) association analysis of DNAm and gene expression, using all individuals with available data (234 AA, 386 HA, and 582 EA individuals), with ESS as the outcome and DNAm as exposure; and (2) association analysis of gene expression (as exposure) and squared-root ESS (as the outcome) in the set of individuals that had both gene expression and ESS measured. Complete details are provided in the Supplementary Material.

## Results

### Study Characteristics

The MESA sample consisted of 619 individuals (53% female) with a mean age of 68 years, with 132 AA, 202 HA, and 285 EA individuals (Table 1). The mean ESS was 6 and 19% were classified as having EDS (ESS>10), which did not vary significantly by ethnicity in this analytical sample (p=0.20) (although ESS was higher in AA than whites in the larger MESA cohort ^23^). Sleep duration was significantly shorter in the AA group compared to the other groups (5.02×10^-7^). Median AHI was 14.1, and 46% of the sample had moderate or more severe sleep apnea (AHI>15). Across groups, sleep apnea was more prevalent and severe in the HA group (Table 1). Significant differences were observed for BMI, having a college education, smoking status, sleep apnea, sleep duration, dietary long chain score, and Total AHEI and alcohol consumption by ancestry group (Table 1).

**Table 1:**
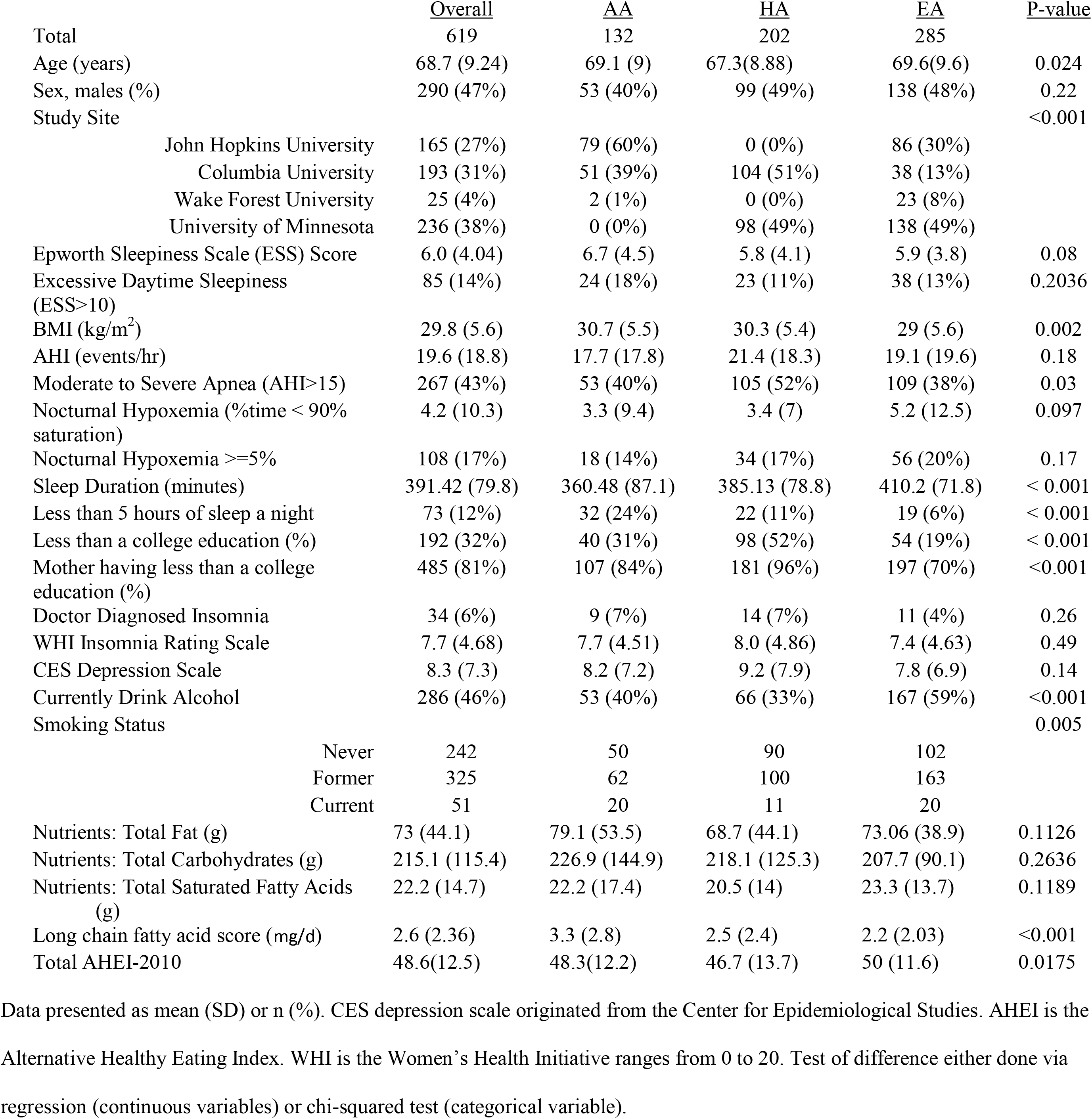
Characteristics of 619 MESA participants. Columns show phenotype characteristics in overall group and each reported ethnicity along with a test for difference in the phenotype between the reported ethnicity.

**Table 2:** Characteristics of 483 CHS participants. Columns show phenotype characteristics as well as separately in European and African Americans.

|  | <u>Overall</u> | <u>African</u><br><u>Americans</u> | <u>European</u><br><u>Americans</u> | <u>P-value</u> |
| --- | --- | --- | --- | --- |
| Total | 483 | 238 | 245 |  |
| Age, years | 74.03 (5.3) | 72.79 (5.4) | 75.23(4.9) | <0.001 |
| Sex, males (%) | 177 (37%) | 82 (34%) | 95 (38%) | 0.3729 |
| Epworth Sleepiness Scale Score | 5.98 (4.1) | 6.38 (4.4) | 5.58 (3.8) | 0.037 |
| Excessive Daytime Sleepiness (ESS>10) | 72 (15%) | 43 (18%) | 29 (12%) | 0.07077 |
| BMI (kg/m<sup>2</sup>) | 27.8 (5) | 29.0 (4.9) | 26.7 (4.8) | <0.001 |
| Currently Drink Alcohol | 209 (43%) | 97 (41%) | 112 (46%) | 0.3136 |
| Smoking Status |  |  |  | 0.07077 |
| Never | 216 (45%) | 108 (45%) | 108 (44%) |  |
| Former | 207 (43%) | 93 (39%) | 114 (47%) |  |
| Current | 60 (12%) | 37 (16%) | 23 (9%) |  |

Data is presented as mean (SD) or n (%).

### Associations Between Sleepiness and DNAm

In the combined analysis across race/ethnic groups, no significant associations were observed between DNAm M-values and ESS. In race/ethnicity stratified analyses, significant associations were only observed in the AA group, where 61 DNAm probes were significantly associated with ESS at q-value ≤ 0.1 (Table 3). The top probe was in *KCTD5* (p=3.96 × 10^-7^), a gene with known sleep-wake effects^58^. Two significant probes were within *RAI1*, a gene associated with several sleep and circadian phenotypes^59,60^, and 5 probes were within *KIF3*, involved in cytokinesis with effects on photoreceptors^61^. We display the top three associations in Figure 2. In 56 of the 61 probes, decreased methylation was associated with more sleepiness (i.e., higher ESS scores) (Supplement Table 2). Testing for any interaction effects between these 61 probes and selfreported ethnicity on the transformed ESS scores in the overall group identified 58 of the 61 associations with a p-value for interaction less than 0.05. Table 3 provides all associations passing the significance threshold, along with their corresponding genes (from Illumina annotation), and replication p-values in CHS. Supplemental Table 2 includes additional information on these associations, including interaction p-values, and effects and standard errors in the overall group, HA, AA and EA, as well as estimated effect sizes, SDs, replication p-values (one-sided) and usual two-sided p-values in the CHS replication samples. Histogram of p-values in each analysis are provided in the Supplementary Material.

**Figure 2:**
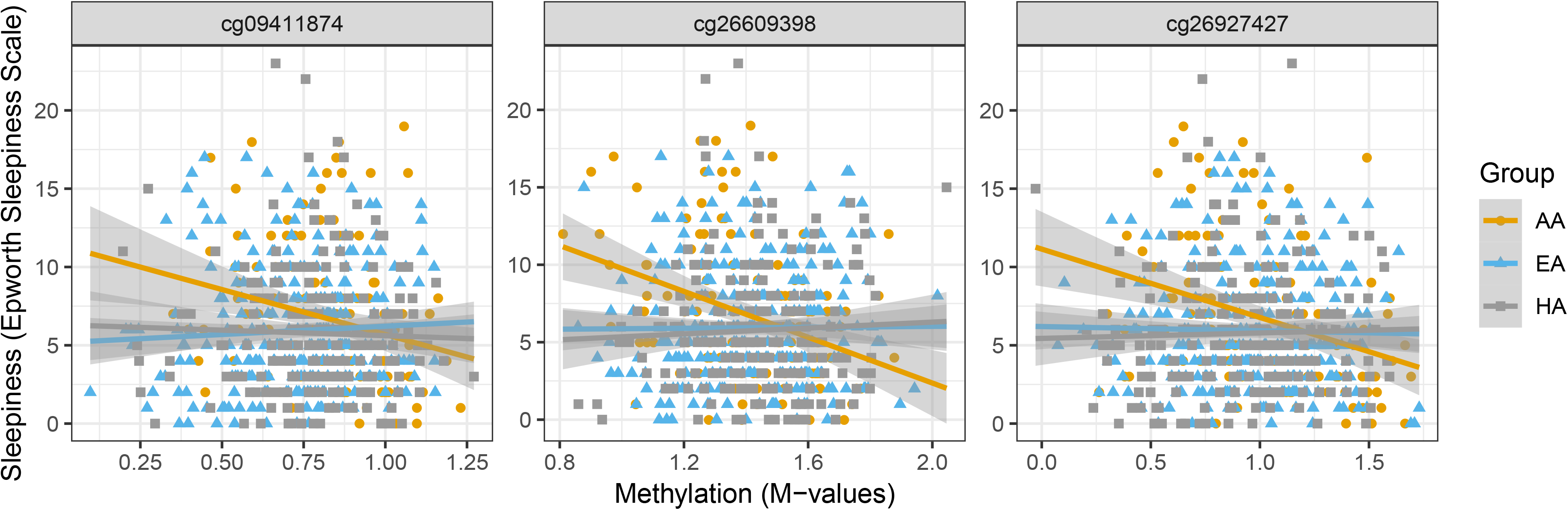
Top three association in AA. Left column shows the probe vs squared-root transformed ESS in the overall sample, right column shows the probe vs transformed ESS in AA only.

**Table 3:**
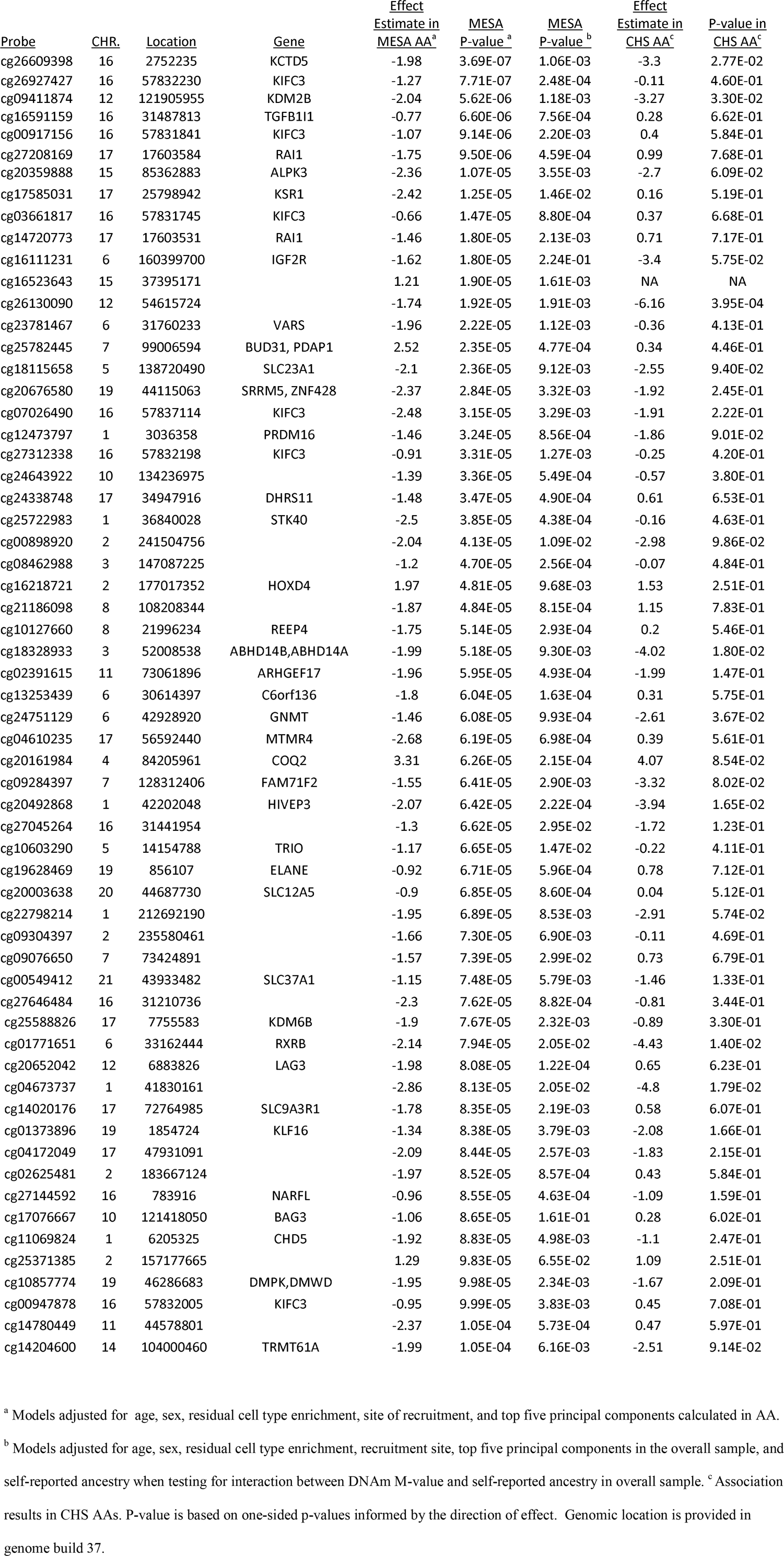
Association analysis results in the MESA AA discovery sample and in the CHS AA replication sample. Table rows correspond to methylation sites associated with the Epworth Sleepiness Scale (square root transformed) with FDR<10% in the discovery sample.

### Sensitivity Analysis

We next evaluated potential explanatory factors or confounders between the ESS-associated DNAm probes in the AA sample. Supplement Table 3 shows p-value between all considered sensitivity parameters and partial R^2^ with ESS after adjusting for age and sex. WHI insomnia score, alcohol use, mother having less than a college education, and sleep duration were all associated with sleepiness (square-root transformed ESS) at a p-value of 0.1 adjusting for age and sex in the AA group (Supplementary Table 3). We included these four variables as additional covariates in the primary model to assess if they modified the association between DNAm and ESS. In our 61 probes, we see a change in the effect size over 10% for 45 of the probes (Supplement Table 4). However, the change for any individual variable was less than 10% (Supplement Table 4). We additionally performed a permutation study to compare the observed changes to what would be expected by chance. The change in association of 51 DNAm sites was significantly different from zero with permutation p-value<0.05, i.e., with less than 5% of permutations demonstrating the same or higher change in estimated association (Supplement Table 4 last column). Supplement Table 4 summarizes the results for the top ten probes. This suggests that the selected exposure variables confound/modify the DNAm-ESS association. Supplemental Table 4 also shows how much the association between DNAm and transformed ESS change after the inclusion of each individual sensitivity parameter by itself. There is some evidence of confounding (effect change over 10% for 45 of the 61 probes). The majority of adjusted associations are still nominally significant (range 2.33 × 10^-3^ to 4.25 × 10^-6^) suggesting that these variables do not explain away the association between DNAm and ESS.

### Replication of DNAm-sleepiness association in CHS

Of the 61 probes found significant in our discovery analysis, 60 were assayed in the CHS AA dataset (n=238). Replication analysis results are provided in Table 3. Globally, the direction of DNAm-sleepiness associations largely agreed between MESA and CHS AA for these 61 probes (binomial p-value=0.005, Supplement Table 2). Forty of the sixty (67% agreement) probes had the same effect sign. Only one probe of the 60 was significant at a Bonferroni level of 0.05 in the CHS AA analysis, cg26130090 (32945 base pairs from *SMUG1* chromosome 12 (MESA p-value = 1.92 ×10^-5^, CHS p-value=3.95 ×10^-4^)). The CHS EA analysis (n=245) agreed with the MESA EA results, which showed no evidence of association in these methylation sites. Consistent with null results in MESA EA, no individual associations replicated in CHS EA, and globally, there was no agreement in the directions of associations between MESA AA and CHS EA (binomial p-value = 0.9393). There was no agreement in direction of effect comparing MESA AA vs MESA EA or MESA HA.

### Cis-meQTL analysis and UK Biobank replication

We performed cis-meQTL analysis for the top ten probes which mapped to seven genes that harbored significant DNAm-sleepiness associations in our AA sample. Thirty SNPs in four genes (*KIFC3, KDM2B, TGFB1L1*, and *RAI1*) passed screening step 1 (P<0.005; Supplementary Table 5). Of these, 15 SNPs passed screening step 2 and were carried forward for testing the SNP-sleepiness association in the UK Biobank data set of European Ancestry individuals. We determined significance in this replication by a p-value threshold of 0.05/15 = 0.003. The results from this analysis are reported in Supplement Table 6. A single SNP passed this threshold, rs9896285 in *RAI1* (p=0.0013).

### Gene expression analysis

Comprehensive results are provided in Supplementary Tables 7 and 8. Briefly, we considered three genes corresponding to the top 10 DNAm sites: *KCTD5* (two transcripts, one DNAm site), *KDM2B* (one transcript and one DNAm site), and *RAI1* (two transcripts and two DNAm sites). For *KDM2B*, cg09411874 methylation was associated with increased expression in the HA group (p=0.04), but there was no strong evidence of *KDM2B* expression association with ESS (p=0.18 in the HA group). The two DNAm sites associated with *RAI1* had p=0.08 in the combined cohort in association with expression of one of the *RAI1* transcripts. However, the estimated association of this transcript with ESS was low (and with p>0.6 for all analyzed groups). Interestingly, larger expression of the other transcript of *RAI1* was significantly associated with decreased ESS (p=0.009 in the combined cohort). We did not detect any associations in the analysis of *KCTD5.*

## Discussion

This study reports for the first time a large-scale DNA methylation association analysis in a multi-ethnic population designed to detect DNAm markers associated with excessive daytime sleepiness. We analyzed the association both in the pooled MESA cohort and separately in each race/ethnicity group (AA, EA, and HA). In our initial discovery stage, there were 61 DNAm markers associated with sleepiness at the FDR rate of < 0.1, solely in the AA group. Of these, 60 were studied in association with the ESS score in the CHS cohort. One of the associations, that of cg26130090 (intergenic), replicated in CHS AAs (p-value =0.0004), and there was an overall directional agreement in the significant associations in MESA AA and CHS AA (Binomial p-value=0.005). Additional top DNAm associations were observed in the *RAI1* and *KCTD5* genes, which have strong biological relevance and were previously reported as associated with several sleep traits^58-60,62-64^.

Our study focused on the multi-ethnic epigenetic bases of daytime sleepiness. Several studies have shown that even after accounting for sleep duration and sleep disorders, sleepiness is less in EA compared to other ethnic groups^23,65-68^. Compared to EA groups, AAs also have a higher prevalence of both short and long sleep durations and sleep apnea syndrome (defined by the presence of an elevated apnea hypopnea index plus daytime sleepiness^23^) and in the larger MESA cohort, AA had higher ESS than EA^23^. Race/ethnicity differences in sleep patterns and susceptibility to EDS appear early in life. AA children have been shown to have shorter sleep duration than white children as early as one year of age^69^. In children with sleep apnea treated with adenotonsillectomy, AA children have higher sleepiness scores than EA children both at baseline and following treatment^70^. Here, in our study of a middle-aged to older community multi-racial sample, we found significant epigenetic findings only in the AA sample. The specificity for the findings in the AA sample could reflect differences in underlying genetic architecture that influence susceptibility to methylation or differences in environmental exposures, such as air pollution, or stress that differentially affect different groups^71-74^. We explored this latter hypothesis by a series of sensitivity analyses that explored potential explanatory effects due to sleep variation and other lifestyle exposures. These analyses showed that adjusting for mother having less than a college education, alcohol use, sleep duration, and insomnia modestly attenuated the effect estimates, suggesting potential mechanisms for epigenetic findings. However, these measures do not characterize lifetime exposures nor included precise measurements of the social or physical environments, emphasizing the need for additional research that links key aspects of an individual’s environmental and sociological exposures with epigenetic markers and health conditions such as sleepiness as important routes for understanding health disparities.

A few of our sleepiness associated DNAm sites are within genes that have strong evidence for associations with sleep related processes. Our most significant DNAm association with excessive sleepiness (MESA AA p-value 3.69×10^-7^), was for cg26609398 in the *KCTD5* gene. While this probe did not replicate in CHS, it was suggestive (CHS AA p-value 0.028) with the association in the same direction. *KCTD5* is a mammalian ortholog of the Drosophila gene *Inc*, a gene expressed in multiple sleep associated regions in the central nervous system and shown to play a fundamental role in sleep in Drosophila by interacting with Cullin-3 ubiquitin ligase (*Cul3*). When mutated, both genes (*KCTD5* and *Cul3*) produce similar phenotypes characterized by markedly shortened and fragmented sleep together with altered synaptic structure and impaired synaptic transmission in relevant neuronal circuits^58^. This suggests that alterations in protein dynamics (e.g. ubiquitination) might disrupt synaptic function and impact sleep. Future research is needed to examine how epigenetic changes in *KCTD5* and similar genes influence gene expression and core elements of the sleep homeostat, with initial work indicating potential protein interactions with the circadian genes *FBXW11* and *BTRC*^75^.

Two of our top ten associated methylation sites were within *RAI1* (retinoic acid induced-1). *RAI1* encodes a protein that is highly expressed in neuronal tissues and is involved in early neural differentiation and transcriptional regulation of circadian clock components^60,62-64^. Variants in this gene have been associated with circadian rhythm disruption, several abnormal sleep traits, neurobehavioral problems and obesity^62,64^. In addition, SNPs in *RAI1* have been found to be associated with sleep apnea^59^. Moreover, our cis-meQTL analysis detected a SNP in *RAI1* influencing sleepiness in an EA population from the UK Biobank. Our analysis in MESA suggests that the SNP rs9896285 has both direct and indirect (through DNAm) effect on excessive sleepiness. However, the methylation association with *RAI1* did not replicate in CHS. In another secondary analysis, we found that the expression of *RAI1* in mononuclear cells was associated with ESS in MESA. More work is needed to further dissect the association of *RAI1* markers with ESS.

Other findings in our study, while not as strong as *KCTD5* and *RAI1*, have some evidence supporting the epigenetic association between sleepiness and DNAm. A significantly associated DNAm site mapped to *KDM2B*, a histone demethylase gene. DNAm at this gene has been found to be associated with BMI^28,76^. Obesity may promote sleepiness through effects on sleep disruption or the effects of inflammatory cytokines (that are augmented by obesity-associated inflammation) on central processes regulating sleep and sleepiness^77,78^. However, our sensitivity analyses did not support a significant influence of obesity on our observed associations. This could be because BMI does not optimally capture the underlying obesity-related phenotypes, or because methylation of *KDM2B* regulates both BMI and sleepiness via a shared process. The methylation site, cg17585031, was near *KSR1* (Kinase Suppressor of Ras 1), a protein suppressor of RAS signaling, a pathway that controls cellular growth and differentiation^79^. *KSR1* has been found to be up-regulated after sleep-restriction^80^ in 26 participants studied in a laboratory-based assessment. DNAm at this gene has been associated with sleep apnea as well, though in a small study at a high significance level (FDR<0.5)^81^. This *KSR1* probe (cg17585031) did not replicate in CHS AA and had opposite sign of association compared to the probe reported in this gene from the sleep apnea study^81^. A significant association was observed for the CpG site near a master cytokine *TGFB1*. Experimental sleep fragmentations in mice has shown rapid increases in expression of *TGFB1* in the hypothalamus and hippocampus^82^.

A unique feature of this study is study of a multi-ethnic population. We identified DNAm-EDS associations that are only present in AAs. DNAm differences in these sites may explain some of the differences in EDS between AAs and EAs, highlighting potential mechanisms to explore in studies of health disparities. The main limitation of our association study is that we had small sample sizes in the group-specific analyses. Despite the limited sample size, we could directly replicate one association (after accounting for multiple testing) and observed a global consistency of the significant association between the discovery cohort MESA AA, and the replication study of CHS AAs. Such global consistency was not observed between MESA AA and CHS EA, strengthening the race/ethnic specific association evidence. Lack of further replication may relate to many factors, including the older sample of CHS participants (who are 5 years older than MESA participants, on average), biases and misclassification in self-reported sleepiness assessment^83^, and potential difference in their exposure history compared to MESA participants. Another limitation was our pruning of DNA methylation sites was stringent though similar has been done prior ^39^. Given our small sample size, this was a good way to reduce the multiple testing burden by excluding probes that were likely not informative. Finally, a limitation which is shared by many epigenome-wide association studies, is the lack of identified exposure explaining the change in methylation and the group-specificity of the associations. We did study potential mechanism by considering cis-meQTLs analysis and a sensitivity analysis adjusting for various, plausible, exposure variable (sleep, lifestyle, and nutrition variables) which may be independently associated with either DNAm, ESS, or both, but these analyses did not lead to strong hypotheses about causal pathways.

In conclusion, we identified multiple DNAm probes associated with sleepiness. The majority of these probes indicated that hypomethylation was positively associated with sleepiness, with some of our top probes being near genes known to be associated with critical neuronal processes that influence sleep. Taken together, our findings suggest that these sleepiness-associated DNAm sites are related to genes that have a biologically meaningful function with regards to sleep. The significance of our findings in AAs identifies biological areas that may contribute to the observed differences in sleepiness between AAs and EAs, providing an avenue for further investigation, both genetic and environmental, of health disparities. For example, within MESA, the *RAI1* SNP (that associates with cg27208169) had MAF of 0.28 in AA, 0.04 in EA, and 0.11 in HA. In the UK biobank, the MAF was 0.06. As mentioned in the introduction, poor sleep outcomes are associated with negative life outcomes and a recent review presented extensive evidence that obstructive sleep apnea and aging share various biological pathways^84^. Future studies with larger sample sizes may elucidate these genomic pathways and how they relate to potential biological or sociological patterns that are being reflected in excessive sleepiness.

## Acknowledgments

We thank the participants of the UK Biobank; replication of the SNP sleepiness association was performed under application 6818. RTB was funded by NCI grant T32CA094880 and NHLBI grant R01HL113338. SR and TS were partially funded by R35 HL135818. HW was supported by NIH NHLBI R01HL113338 (to SR) and Sleep Research Society Foundation Career Development Award 018-JP-18 (to HW). RS was supported by NIH R01DK107859, R01DK102696 and R01DK105072.

MESA:

This research was supported by the Multi-Ethnic Study of Atherosclerosis (MESA).

MESA and the MESA SHARe project are conducted and supported by the National Heart, Lung, and Blood Institute (NHLBI) in collaboration with MESA investigators. Support for MESA is provided by contracts HHSN268201500003I, N01-HC-95159, N01-HC-95160, N01-HC-95161, N01-HC-95162, N01-HC-95163, N01-HC-95164, N01-HC-95165, N01-HC-95166, N01-HC-95167, N01-HC-95168, N01-HC-95169, UL1-TR-000040, UL1-TR-001079, UL1-TR-001420, UL1-TR-001881, and DK063491.

MESA SLEEP was funded by R01HL098433. Funding support for the Sleep Polysomnography dataset was provided by grant HL56984.

The MESA Epigenomics Studies were funded by R01HL101250, R01 DK103531-01, R01DK103531, R01 AG054474, and R01 HL135009-01 to Wake Forest University Health Sciences.

The provision of genotyping data was supported in part by the National Center for Advancing Translational Sciences, TSCI grant UL1TR001881, and the National Institute of Diabetes and Digestive and Kidney Disease Diabetes Research (DRC) grant DK063491. Funding for MESA SNP Health Association Resource (SHARe) genotyping was provided by NHLBI Contract N02-HL-64278. Genotyping was performed at Affymetrix (Santa Clara, California, USA) and the Broad Institute of Harvard and MIT (Boston, Massachusetts, USA) using the Affymetrix Genome-Wide Human SNP Array 6.0.

The authors thank the other investigators, the staff, and the participants of the MESA study for their valuable contributions. A full list of participating MESA investigators and institutions can be found at http://www.mesa-nhlbi.org.

CHS: Infrastructure for the CHARGE Consortium is supported in part by the National Heart, Lung, and Blood Institute grant R01HL105756. The CHS research was supported by NHLBI contracts HHSN268201200036C, HHSN268200800007C, HHSN268201800001C, N01HC55222, N01HC85079, N01HC85080, N01HC85081, N01HC85082, N01HC85083, N01HC85086; and NHLBI grants U01HL080295, U01HL130114, K08HL116640, R01HL087652, R01HL092111, R01HL103612, R01HL105756, R01HL103612, R01HL111089, R01HL116747 and R01HL120393 with additional contribution from the National Institute of Neurological Disorders and Stroke (NINDS). Additional support was provided through R01AG023629 from the National Institute on Aging (NIA), Merck Foundation / Society of Epidemiologic Research as well as Laughlin Family, Alpha Phi Foundation, and Locke Charitable Foundation. A full list of principal CHS investigators and institutions can be found at CHS-NHLBI.org. The provision of genotyping data was supported in part by the National Center for Advancing Translational Sciences, CTSI grant UL1TR001881, and the National Institute of Diabetes and Digestive and Kidney Disease Diabetes Research Center (DRC) grant DK063491 to the Southern California Diabetes Endocrinology Research Center.

The content is solely the responsibility of the authors and does not necessarily represent the official views of the National Institutes of Health.

## Abbreviations

AAs: African American
AHI: Apnea-Hypopnea Index
ANOVA: Analysis of Variance
BMI: Body mass index
CES: Center for Epidemiological Studies
CHS: Cardiovascular Health Study
DNAm: DNA methylation
EAs: European Americans
EDS: Excessive daytime sleepiness
ESS: Epworth Sleepiness Scale
FDR: False Discovery Rate
GWAS: Genome-Wide association study
HAs: Hispanic Americans
meQTLs: methylation expressive quantitative trait loci
MESA: Multi-ethnic Study of Atherosclerosis
PC: Principal Component
Per90: nocturnal hypoxemia
SNP: Single Nucleotide Polymorphism
WHI: Women’s Health Initiative

